# A novel transcriptional signalling pathway mediated by the trafficking protein Ambra1 via scaffolding Atf2 complexes

**DOI:** 10.1101/2020.01.08.899328

**Authors:** Christina Schoenherr, Adam Byron, Margaret C Frame

## Abstract

Previously, we reported that Ambra1 is a core component of a cytoplasmic trafficking network, acting as a spatial rheostat to control active Src and FAK levels in addition to its critical roles in autophagy during neurogenesis. Here we identify a novel nuclear scaffolding function for Ambra1 that controls gene expression. Ambra1 binds to nuclear pore proteins, to other adaptor proteins like FAK and Akap8 in the nucleus, as well as to chromatin modifiers and transcriptional regulators such as Brg1, Cdk9 and the cAMP-regulated transcription factor Atf2. Ambra1 contributes to their association with chromatin and we identified genes whose transcription is regulated by Ambra1 complexes, likely via histone modifications and phospho-Atf2-dependent transcription. Therefore, Ambra1 scaffolds protein complexes at chromatin, regulating transcriptional signalling in the nucleus; in particular, it recruits chromatin modifiers and transcriptional regulators to control expression of genes such as *angpt1, tgfb2, tgfb3, itga8* and *itgb7* that likely contribute to the role of Ambra1 in cancer cell invasion.

## INTRODUCTION

Ambra1 (Activating Molecule in Beclin1-Regulated Autophagy) is already known to be an important protein in physiology, e.g. in the development of the central nervous system, vertebrate embryogenesis, adult neurogenesis and cancer cell invasion (1–5). However, our understanding of the full range of functions of this crucial cellular regulator is unknown. As an important autophagy regulator, Ambra1 binds Beclin1 and is involved in the initiation of autophagy that is needed for neurogenesis (2). In the absence of autophagy, the Ambra1/Beclin1/Vps34 complex is bound to the dynein motor complex; when autophagy is induced, the kinase ULK1 phosphorylates Ambra1, resulting in its release from the dynein complex (6,7). Additionally, the function of Ambra1 is negatively regulated by mTOR that suppresses its binding to the E3-ligase TRAF1 and the ubiquitylation of ULK1, thereby controlling the stability and function of ULK1 (8). During apoptosis, caspases and calpains mediate cleavage as well as degradation of Ambra1 (9). Furthermore, Ambra1 expression is regulated by RNF2-dependent ubiquitylation, resulting in degradation (10). Ambra1 is also involved in the regulation of mitophagy (11).

Ambra1 has been both positively and negatively implicated in cancer. Thus far, it has been proposed as a tumour suppressor, supporting the binding of c-Myc to the phosphatase PP2A, resulting in c-Myc degradation as well as reduced proliferation and tumourigenesis (12). Ambra1 has been positively implicated in cholangiocarcinoma, where overexpression is correlated with invasion and poor survival (5). In addition, through its ability to bind PP2A, Ambra1 stabilises the transcription factor FOXO3, triggering FOXP3-mediated transcription, and T-cell differentiation and homeostasis (13).

We reported previously that in squamous cell carcinoma (SCC) cells derived from the mutated oncogenic H-Ras driven DMBA/TPA model of carcinogenesis, Ambra1 is a Focal Adhesion Kinase (FAK)- and Src-binding partner, regulating cancer-related phenotypes like cancer cell polarisation and chemotactic invasion (4,14). In FAK-depleted SCC cells, Ambra1 is involved in the targeting of active Src to intracellular autophagic puncta, while an Ambra1-binding impaired FAK mutant retains more active FAK and Src at focal adhesions, resulting in increased cell adhesion and invasion. We concluded that Ambra1 lies at the heart of an intracellular trafficking network in SCC cells, regulating the localisation of active FAK and Src required for cancer processes (4).

Here, we investigate the nuclear function of Ambra1, and show that it binds to FAK in the nucleus, as well as to other nuclear adaptor proteins, nuclear pore components, histone modifying enzymes and regulators of transcription – in some cases regulating their recruitment to chromatin. Specifically, Ambra1 forms complexes with Akap8, Brg1 as well as Atf2 and is responsible for the recruitment of Akap8, Bgr1, the Mediator complex components Cdk9 as well as p-Atf2 T71 to chromatin. Both Ambra1 and its binding scaffold protein Akap8 regulate the binding of transcriptional proteins to chromatin, especially p-Atf2 T71, and modulate histone modifications. The binding of Atf2/p-Atf2 T71 to chromatin is most likely regulated by the Ambra1-interacting proteins Cdk9. Therefore, Ambra1 acts as a scaffold protein in the nucleus, recruiting regulators of transcription to chromatin. This creates an Ambra1-dependent nuclear microdomain that regulates gene expression.

## RESULTS

### Ambra1 localises to the nucleus

Here we show that Ambra1 not only locates at focal adhesions and in intracellular autophagic puncta in SCC cells, but also in the nucleus (Figure 1A). Staining with only secondary antibodies (anti-rabbit 488 and anti-mouse 594) ruled out unspecific false-positive nuclear staining (Supplementary Figure 1A). In order to confirm Ambra1 was nuclear biochemically, fractions of SCC FAK-WT and -/- were isolated and subjected to Western blotting (Figure 1B). Biochemical nuclear isolations were checked by blotting with anti-GAPDH, anti-Lamin A/C and anti-H4. In the nuclear fractions of these SCC cells, we could detect Ambra1 as well as FAK, the latter in line with our previous reports (15,16). The nuclear localisation of Ambra1 was not dependent on FAK, as nuclear Ambra1 is present at indistinguishable levels in nuclear fractions from both FAK-deficient (FAK -/-) SCC cells and the same cells reexpressing wild type FAK to similar levels as parental SCC cells (FAK-WT) (17,18). In more highly purified cellular fractions of SCC FAK-WT and -/- cell lysates, extracting cytosolic (C), perinuclear (PN) and nuclear (N) fractions (Supplementary Methods, Supplementary Figure 1B), Ambra1 was present at comparable levels in the cytosolic and nuclear fractions, and at higher levels in the perinuclear fraction. Fraction purities were confirmed by blotting respective fractions with anti-GM130, anti-PDI, anti-Lamin A/C and anti-GAPDH (Supplementary Figure 1B). Further, in contrast to FAK, Ambra1 can also be detected in nuclear extracts from primary mouse keratinocytes (Supplementary Figure 1C) (16).

**Figure 1:**
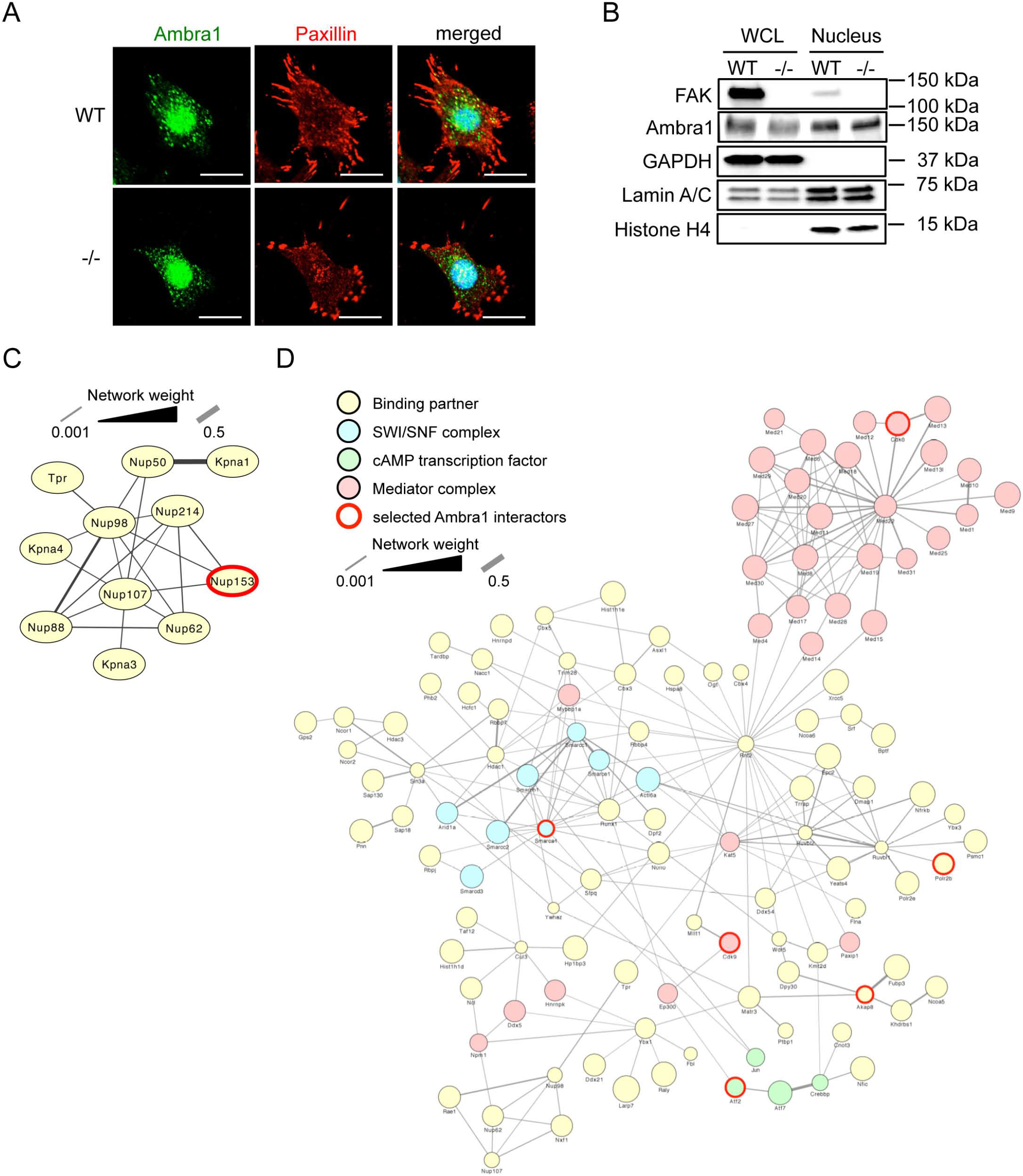
Nuclear Ambra1 binds chromatin modifiers and transcriptional regulators. **(A)** Representative immunofluorescence images of SCC FAK-WT and -/- cells which were grown on glass coverslips for 24 h, fixed and stained with anti-Ambra1, anti-Paxillin and DAPI. Scale bars, 20 μm. **(B)** Whole cell and nuclear lysates of SCC FAK-WT and -/- cells were subjected to Western blot analysis with anti-Ambra1 and anti-FAK. Anti-GAPDH, anti-Lamin A/C and anti-H4 served as controls for the purity of the nuclear extracts as well as loading controls. **(C, D)** Gene ontology enrichment analysis for biological processes of nuclear Ambra1-binding proteins forming part of the nuclear pore complex **(C)** and being involved in transcription **(D)** in SCC FAK-WT and -/- cells. Hits were filtered for statistical significant (p < 0.05) 2-fold enrichment over IgG control and subsequently used to build a protein interaction network based on direct physical interactions (grey lines). Components of various complexes involved in transcription are highlighted: SWI/SNF complex (blue), cAMP regulated transcription factor (green) and Mediator complex (red). Ambra1-interacting proteins selected for further investigation are highlighted with a red circle.

### Nuclear Ambra1 binds proteins involved in transcription

Next, we investigated the nature of protein binding partners of Ambra1 in the nucleus. Highly purified nuclear extracts of SCC FAK-WT and -/- cells were obtained by sucrose gradient centrifugation and used for Ambra1 immunoprecipitations (antirabbit IgG served as a negative control), and specific nuclear binding proteins were determined by quantitative label-free mass spectrometry (see Materials and Methods). Amongst the nuclear Ambra1 interacting proteins identified were several proteins that form part of the nuclear pore complex (functional interaction network shown in Figure 1C). Indeed, we had previously observed that the nuclear pore protein Tpr was an Ambra1-interacting protein using whole cell lysates (4) – together implying that Ambra1 was associated with nuclear pore proteins either during entry into the nucleus or as part of its nuclear functions. Gene ontology analysis for biological processes of proteins that bind Ambra1 in nuclear fractions revealed transcription, RNA/mRNA processing, histone modification and chromatin modification as the most highly represented categories (Supplementary Figure 2). As a result of these analyses, we further examined nuclear Ambra1-interactors involved in the regulation of transcription and used these hits to build a protein interaction network based on known direct physical interactions (Figure 1D). Amongst these were several components of the Mediator complex, a multiprotein complex functioning as a transcriptional coactivator for RNA Polymerase II (highlighted in red) (19,20). Also present were several components of the SWI/SNF (SWItch/Sucrose Non-Fermentable) nucleosome remodelling complex that allows transcription factor binding by ‘opening up’ chromatin structure, e.g. the catalytic subunit SMARCA4 (Brg1), which allows ATP dependent chromatin remodelling (highlighted in blue) (21–23). Nuclear Ambra1 was also found to interact with several members of the cAMP dependent AP-1 complex, including c-Jun, Fosl, Atf2 (a member of the CREB (cAMP response element binding) family of leucine zipper proteins) and Atf7, which also binds to nuclear FAK (highlighted in green; (16,24). Since cAMP-regulated transcription factors were amongst nuclear Ambra1-binding proteins, we also noted that nuclear Ambra1 binds to Akap8 (A kinase anchor protein 8, also known as Akap95), a scaffold that targets PKA to cAMP-responsive elements in the nucleus, and which is linked to chromatin status and retention of p90 S6K to the nucleus (25–28). Interestingly, both Atf2 and Akap8 were also identified by proteomics as Ambra1 binding proteins using whole cell lysates in our previous experiments (4).

We next selected a number of proteins identified by mass spectrometry analyses, i.e. Nup153, Akap8, Brg1, Atf2, and the RNA polymerase II Rpb1 (all highlighted with red circles in Figure 1C and D) and confirmed their binding with Ambra1 in the nucleus by co-immunoprecipitation in nuclear fractions (Figure 2A – E). One question we had, given our previously reported co-functioning of Ambra1 and FAK in the cytoplasm, was whether or not FAK may be regulating the nuclear translocation of its binding partners, such as Ambra1 that also locates to the nucleus. We did not find any significant difference in the nuclear levels of Ambra1 or its interaction with the nuclear binding partners examined between SCC cells expressing FAK-WT when compared to FAK-deficient (-/-) cells; we therefore conclude that FAK does not regulate the trafficking of Ambra1 to the nucleus or Ambra1 associations there. Therefore, in future experiments, we have generally only presented data from SCC cells expressing FAK-WT.

**Figure 2:**
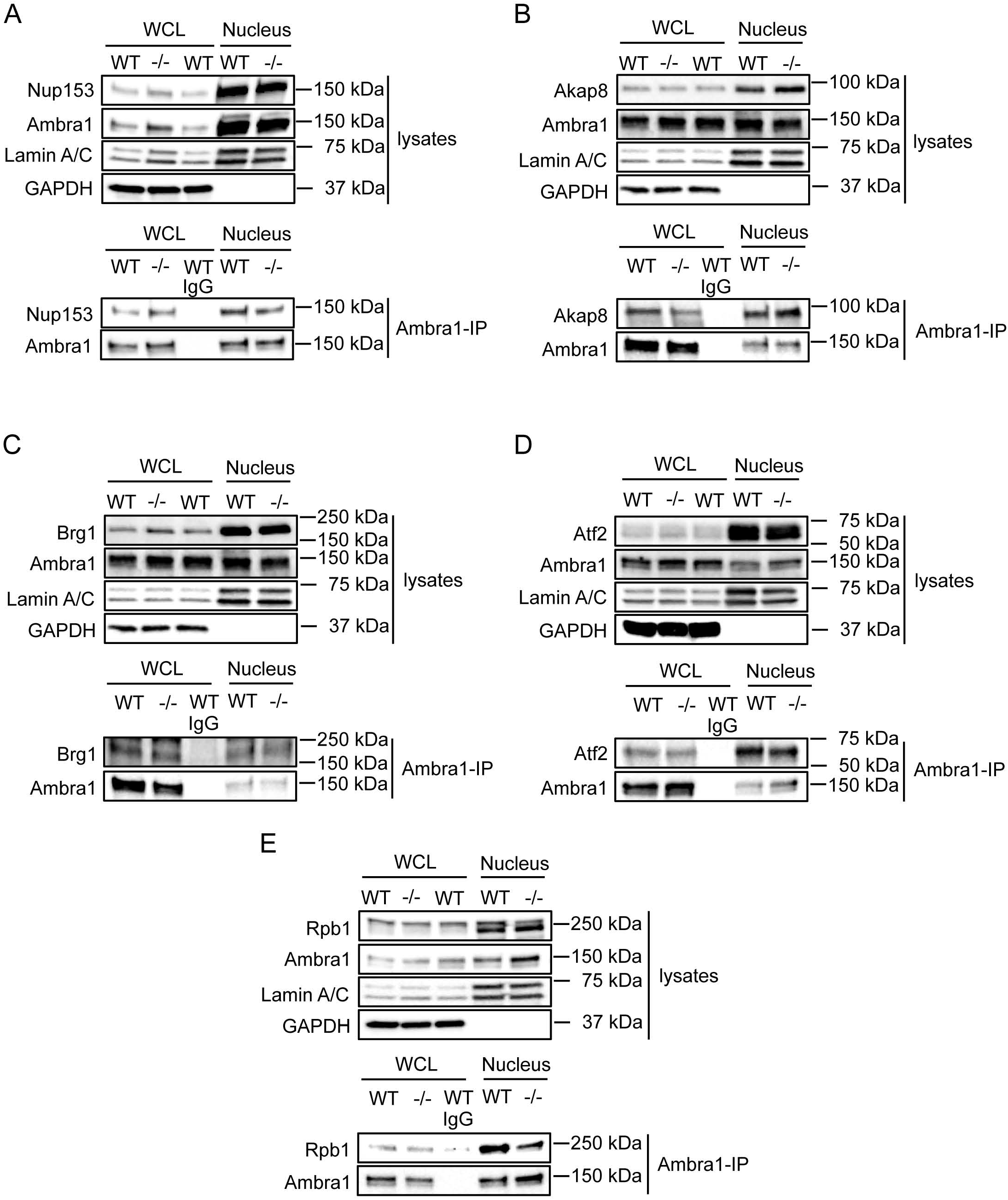
Nuclear Ambra1 binding to mass spectrometry-identified interaction partners. Ambra1 was immunoprecipitated from whole cell and nuclear lysates of SCC FAK-WT and -/- cells using anti-Ambra1, followed by Western blot analysis with anti-Ambra1 and anti-Nup153 **(A)**, anti-Akap8 **(B)**, anti-Brg1 **(C)**, anti-Atf2 **(D)** and anti-Rpb1 **(E)**. Anti-Lamin A/C and anti-GAPDH were used as a control for the purity of the nuclear lysates as well as a loading control.

### Loss of Ambra1 causes reduced association of interacting partners with chromatin

Since Ambra1 has predominantly been defined as a scaffold protein involved in intracellular trafficking/autophagy thus far, we hypothesized that Ambra1 might also serve as a scaffold protein in the nucleus. Therefore, we investigated whether reducing Ambra1 levels using efficient siRNA pool-mediated depletion that we have used previously (4) did not significantly alter the levels of its interacting proteins in whole cell lysates or nuclear extracts (Figure 3A). We next isolated chromatin from SCC FAK-WT cells after Ambra1 depletion and probed for FAK, Brg1, Akap8, Atf2 and its phosphorylated (and activated) form p-Atf2 T71 (Figure 3B, C). Reduced expression of Ambra1 suppressed the binding of FAK, Brg1, Akap8 and p-Atf2 (but not visibly total Atf2) to chromatin to a greater or lesser extent. This suggests that, as well as Ambra1-mediated chromatin recruitment, there are likely other routes by which binding partners are recruited to chromatin since their recruitment is reduced, but not diminished, upon Ambra1 loss. In these analyses, we also included the Cdk8 and Cdk9, components of the Mediator complex that modulates RNA polymerase II-mediated transcription (29–31). These were included because they were: (1) identified as binding to Ambra1 in mass spectrometry experiments (Figure 1D, highlighted with a red circles), and (2) are predicted to be enzymes with the potential to phosphorylate Atf2 at T71 residue on chromatin that we observe in the presence of Ambra1 (Figure 3; predictions via the Biocuckoo phosphorylation prediction site: Cdk8 (score 40.329; cut-off 29.727), Cdk9 (score 7.393; cut-off 2.931). In keeping with other Ambra1-binding proteins studied here, we found that Cdk9, but not Cdk8, binding to chromatin was reduced upon depletion of Ambra1 (Figure 3B, C). In this regard, we note that Cdk9 has been reported to bind another SWI/SNF complex component, SMARCB1, which has also been identified as a nuclear Ambra1 binding protein (32), implying there are likely other components of Ambra1 complexes that link to chromatin remodelling. Taken together, our data imply that Ambra1 forms complexes in the nucleus with other protein scaffolds, such as the PKA scaffold Akap8, chromatin modifiers and transcription factors, including Atf2.

**Figure 3:**
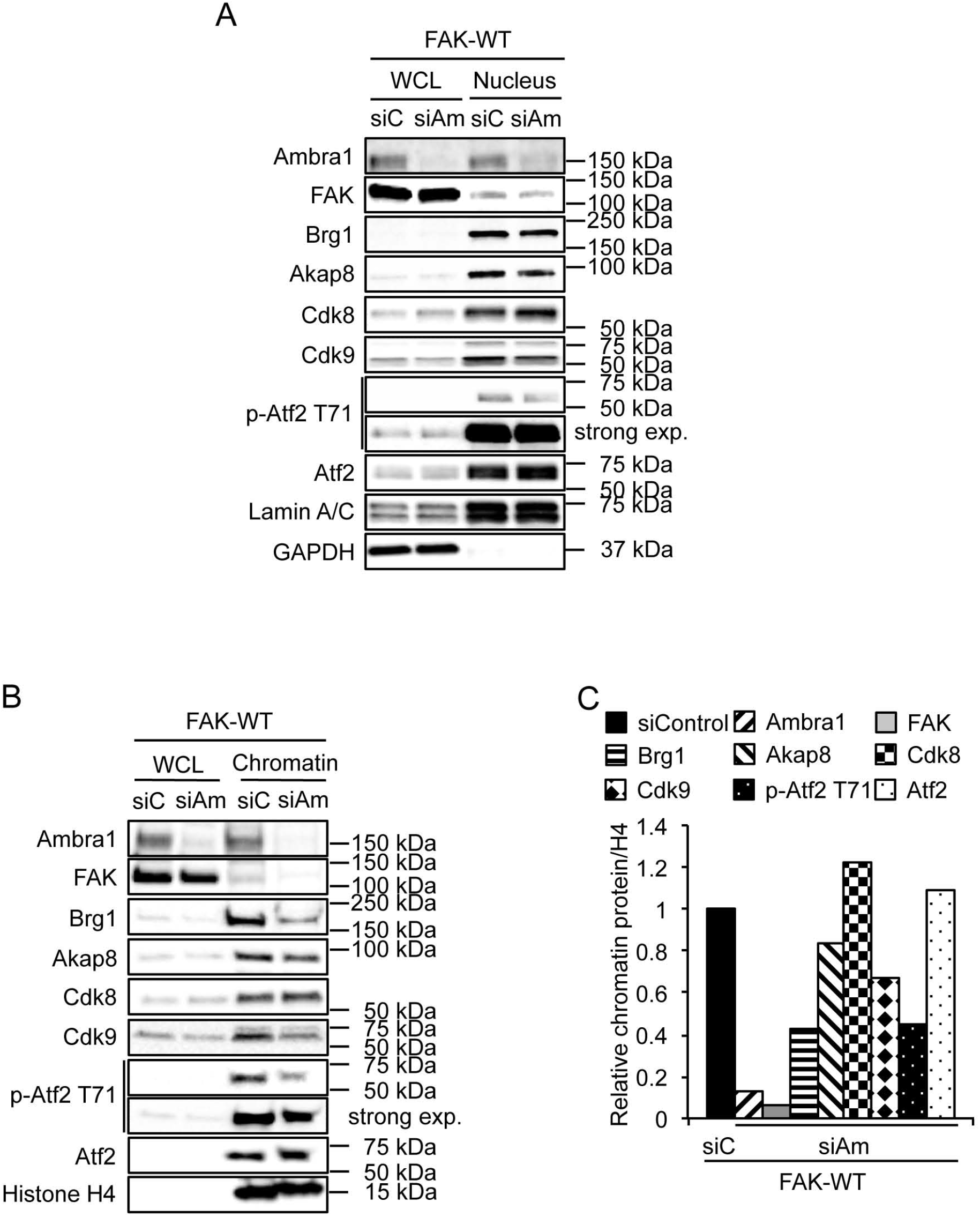
Loss of Ambra1 leads to reduced association of interacting proteins with chromatin. **(A)** SCC FAK-WT cells were transfected with siControl and siAmbra1 (siGENOME pool). After 48 h whole cell and nuclear lysates were analysed by Western blot using anti-Ambra1, anti-FAK, anti-Brg1, anti-Akap8, anti-Cdk8, anti-Cdk9, anti-p-Atf2 T71 and anti-Atf2. Anti-Lamin A/C and anti-GAPDH were used as a control for the purity of the nuclear lysates as well as a loading control. **(B)** SCC FAK-WT cells were transfected with siControl and siAmbra1. After 48 h whole cell lysates and chromatin extracts were analysed by Western blot using anti-Ambra1, anti-FAK, anti-Brg1, anti-Akap8, anti-Cdk8, anti-Cdk9, anti-p-Atf2 T71 and anti-Atf2. Anti-Histone H4 served as a marker for chromatin as well as a loading control. **(C)** The graph shows relative chromatin protein levels normalised to Histone H4.

### Akap8 also regulates the level of p-Atf2 at chromatin

Ambra1 binds the PKA scaffold Akap8 and is required for its optimal binding to chromatin. Upon depletion of Akap8 by pooled siRNA, we found that whilst neither Ambra1 (nor FAK) binding to chromatin was affected, showing it was downstream of Ambra1, the level of p-Atf2 T71 associated with chromatin was reduced (Figure 4A, B). This implies a model (Figure 4E) whereby Ambra1 is upstream of recruitment of Akap8 to chromatin, and Akap8, in turn, is required for optimal chromatin association of active p-Atf2 that is also Ambra1-dependent. Total Atf2 recruitment was not reduced by depletion of Akap8, suggesting that a specific function of Akap8 may be to recruit the enzyme that phosphorylates Atf2 at chromatin. We noted that the activity of Atf2 is proposed to be regulated by phosphorylation at several residues, including T71, by kinases such as ERK, JNK, p38 and PLK3, promoting Atf2 heterodimerisation and increased transcription and histone acetyl transferase (HAT) activity (24,33–39). However, using inhibitors of the kinases proposed above to phosphorylate Atf2 on T71, we did not find evidence for a role for any of these in regulating Atf2 phosphorylation at chromatin in SCC cells used here (not shown). Therefore, we examined the Mediator complex kinases Cdk8 and Cdk9, which, as mentioned previously, may potentially phosphorylate Atf2 at T71. We therefore depleted Cdk8 and Cdk9 in SCC FAK-WT cells and prepared nuclear and chromatin fractions to probe for p-Atf2 T71. We found that depletion of Cdk9 resulted in reduced chromatin-associated p-Atf2 T71 (Figure 4C, D) that is also Ambra1 and Akap8-dependent; however, in this case total Atf2 recruited to chromatin was also reduced. These findings imply that the Mediator complex component, Cdk9, which binds to Ambra1 in the nucleus, controls recruitment of Atf2 to chromatin downstream of Ambra1 and also Atf2 phosphorylation/activation.

**Figure 4:**
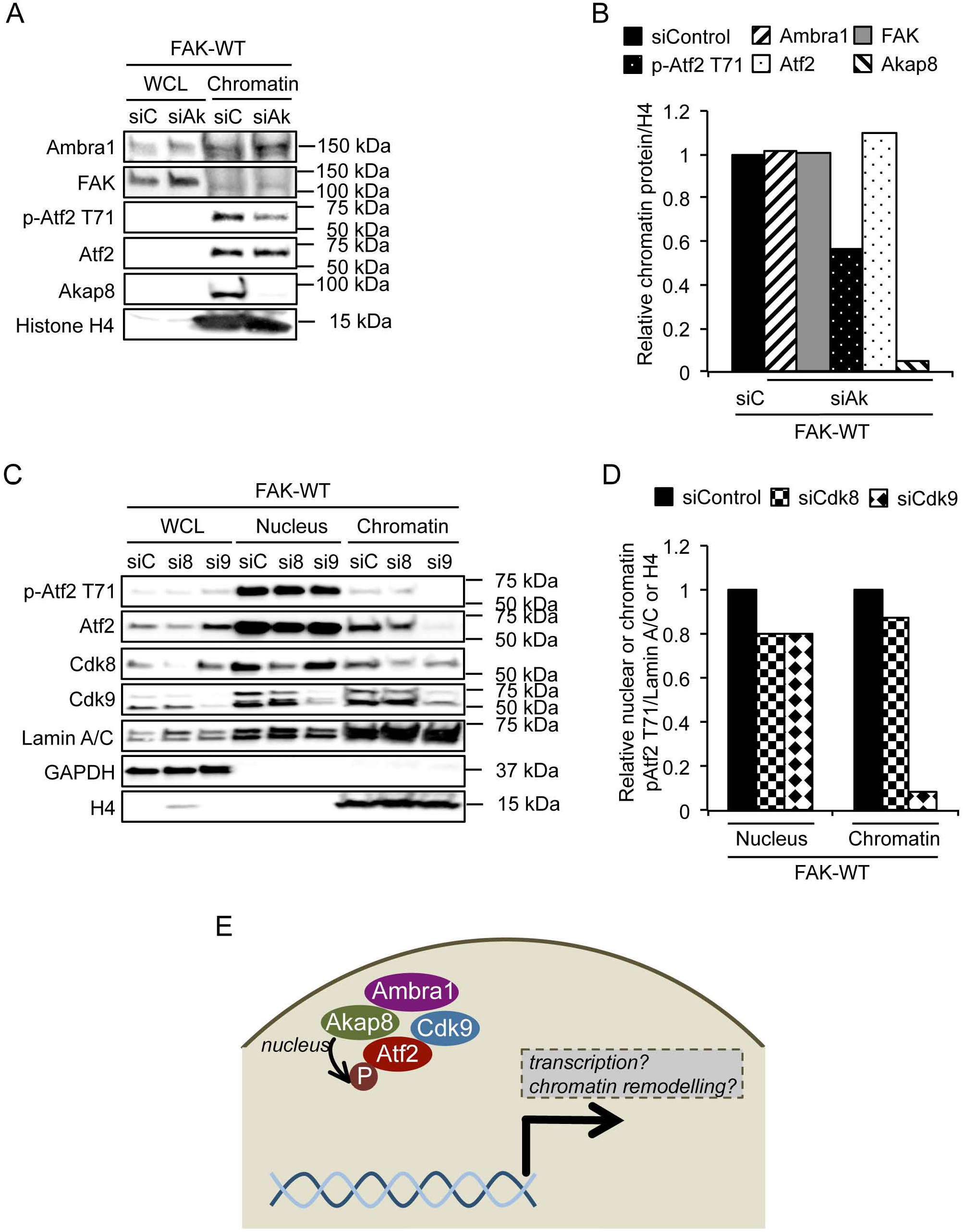
Depletion of Akap8 and Cdk9 reduce p-Atf2 T71 binding to chromatin. **(A)** SCC FAK-WT cells were transfected with siControl and siAkap8 (siGENOME pool). After 48 h whole cell lysates and chromatin extracts were analysed by Western blot using anti-Ambra1, anti-FAK, anti-p-Atf2 T71, anti-Atf2 and anti-Akap8. Anti-Histone H4 served as a marker for chromatin as well as a loading control. **(B)** The graph shows relative chromatin protein levels normalised to Histone H4. **(C)** SCC FAK-WT cells were transfected with siControl, siCdk8 and siCdk9 (siGENOME pool). After 48 h whole cell lysates, nuclear and chromatin extracts were analysed by Western blot using anti-pAtf2 T71, anti-Atf2, anti-Cdk8 and anti-Cdk9. Anti-Lamin A/C, anti-GAPDH and anti-Histone H4 served as a control for the purity of nuclear and chromatin extracts as well as a loading control. **(D)** The graph shows relative nuclear or chromatin protein levels normalised to Lamin A/C or Histone H4 respectively. **(E)** Hypothesis model. In the nucleus of SCC cells Ambra1 and Akap8 form a complex and contribute to the recruitment of active Atf2 (p-Atf2 T71) to chromatin, most likely downstream via the Mediator complex component Cdk9. This strongly suggests that Ambra1 might be involved in chromatin remodeling and transcription. Further, together with Akap8 and Atf2 it might co-regulate the expression of a sub-set of genes.

Thus, Ambra1 and Akap8, which form a complex in the nucleus of SCC cells, both contribute to the recruitment of active Atf2 (p-Atf2 T71) to chromatin, likely via the Mediator complex component Cdk9 downstream (see model in Figure 4E). Therefore, an obvious question that follows is whether Ambra1, Akap8 and Atf2 co-regulate the expression of a subset of genes.

### Ambra1, Akap8, CDK9 and Atf2 coregulate a sub-set of genes

The data presented to this point showed that Ambra1 localises to the nucleus, associates with chromatin and interacts with nuclear proteins that regulate transcription. Both Ambra1 and its binding partner Akap8 recruit transcription factors, such as Atf2; the latter is proposed to result in histone modifications and altered chromatin accessibility, leading to transcriptional changes (40). To address whether there were genes whose transcription was co-regulated by Ambra1, Akap8 and Atf2, SCC FAK-WT cells were transfected with siControl, siAmbra1, siAkap8 or siAtf2 siRNA. A sub-set of genes whose expression was changed by all three depletions was identified using the Nanostring PanCancer Mouse Pathways Panel. In total we identified 94 genes that were significantly (p < 0.05) at least 2-fold up- or downregulated compared to control siRNA (Figure 5A, B; Supplementary Figure 3A – F). Ambra1, Atf2 or Akap8 depletion significantly altered the expression of 18 genes from this panel (Figure 5A, B). In order to validate the gene expression changes, we performed qRT-PCR for the co-regulated genes *angpt1, tgfb2, tgfb3, itga8* and *itgb7* Figure 5d, e). For *angpt1, tgfb2, tgfb3* and *itga8* we confirmed the downregulation upon siRNA transfection (Figure 5E), whilst upregulation of *itgb7* was also confirmed (Figure 5E). In addition, KEGG (Kyoto Encyclopedia of Genes and Genomes) analysis of Ambra1, Akap8 and Atf2 regulated genes revealed the top enriched signalling pathway gene sets ‘PI3K-Akt signalling pathway’, ‘pathways in cancer’, ‘focal adhesion’ and ‘MAPK signalling pathway’ were in common (Figure 5C; for full list see Supplementary Figure 4), strongly suggesting an overlap in the functions of the genes regulated by Ambra1, Akap8 and Atf2 complexes.

**Figure 5:**
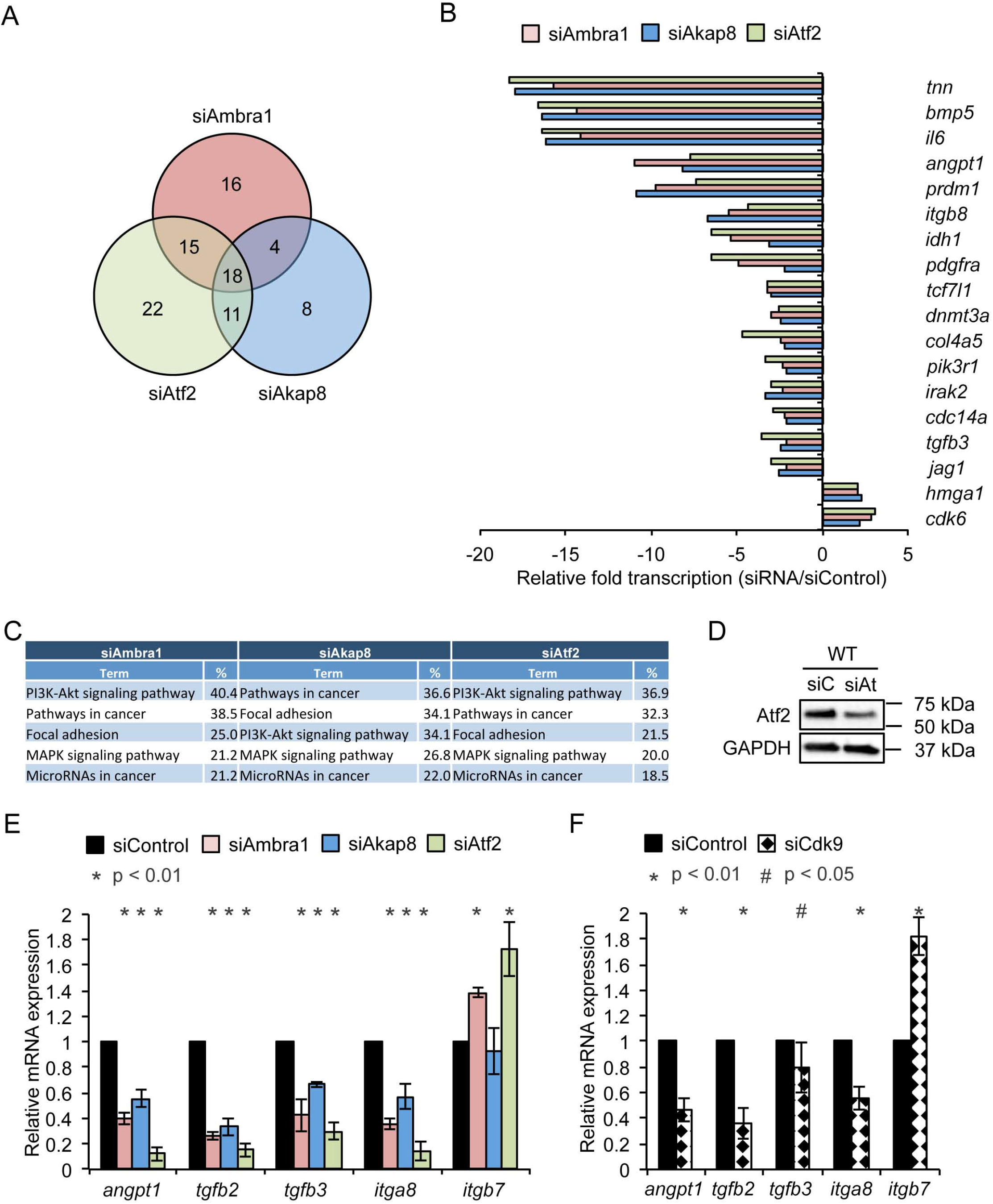
Ambra1, Akap8, Cdk9 and Atf2 co-regulate a sub-set of genes. SCC FAK-WT cells were transfected with siControl, siAmbra1, siAkap8 and siAtf2 (siGENOME pool). RNA was isolated 48 h post transfection and subjected to gene expression analysis using the PanCancer Mouse Pathways Panel. **(A)** Venn diagram of significantly altered genes compared to siControl (p < 0.05, 2-fold difference compared to siControl). **(B)** Relative gene expression of 18 co-regulated genes by Ambra1, Akap8 and Atf2 knockdown compared to siControl. **(C)** Table of the top five enriched signalling pathway gene sets according to KEGG (Kyoto Encyclopedia of Genes and Genomes) pathway analysis of Ambra1, Akap8 and Atf2 regulated genes. **(D)** Western blot showing Atf2 knockdown by siRNA in SCC FAK-WT cells. Anti-GAPDH served as a loading control. **(E, F)** Validation of Ambra1, Akap8 and Atf2 mediated *angpt1, tgfb2, tgfb3, itga8* and *itgb7* expression changes by qRT-PCR using RNA isolated 48 h post transfection of SCC FAK-WT cells with siControl, siAmbra1, siAkap8 and siAtf2 **(E)** as well as siControl and siCdk9 **(F)**. Error bars: s.d. (*) p < 0.01. (#) p < 0.05.

Since we found the nuclear Ambra1-binding Mediator complex protein Cdk9 to be part of the regulation of p-Atf2 at chromatin in SCC cells, (Figure 4C, D), we next addressed whether Cdk9 was also implicated in the transcriptional regulation of the above co-regulated genes. Depletion of Cdk9 by siRNA resulted in broadly similar gene expression changes to that observed upon depletion of Ambra1, Akap8 or Atf2, e.g. expression of *angpt1, tgfb2, tgfb3* and *itga8* was reduced (Figure 5F), whilst *itgb7* was increased in qualitative agreement with the effects of depleting Ambra1, Akap8 or Atf2 (Figure 5F).

Taken together, these results indicate that Ambra1, Akap8, Cdk9 and Atf2 are complexed together and are at chromatin, likely at the promoters of co-regulated genes described above, and potentially many more not represented in the Nanostring PanCancer Mouse Pathways Panel used here. The presence of chromatin modifiers in the nuclear Ambra1 interactome, validated in the case of Brg1 (Figure 1 and 2C), suggested the intriguing possibility that the novel transcriptional regulatory pathway we describe is, at least in part, regulated by chromatin accessibility. In keeping with this, the functioning of Atf2 that is phosphorylated at T71 downstream of Ambra1 and Akap8, is known to promote HAT activity and transcription (35,38).

### Ambra1 and Akap8 regulate histone modifications

Functional interaction network analysis of nuclear Ambra1 binding partners identified by mass spectrometry revealed several components of histone modification complexes, including histone acetylation complexes NSL (Non-specific lethal complex), NuA4 (nucleosome acetyltransferase of H4) and PCAF (p300/CBP-associated factor), as well as histone methylation complexes MLL1/MLL (mixed-lineage leukemia 1) and SET1 (Figure 6A). Further, gene ontology analysis for biological processes of nuclear Ambra1 interactors identified covalent chromatin and histone modifications as well as chromatin remodelling as categories (Supplementary Figure 2). In addition, Akap8 has been reported to bind to Dpy30, a core subunit of H3K4 histone methyltransferases (41). Therefore, we next addressed whether Ambra1 and Akap8 influence histone modifications by transfecting SCC FAK-WT and -/- cells with siControl, siAmbra1 (Figure 6B, C) or siAkap8 (Figure 6D, E) and examined the histone modifications H3K4me2, H3Kme3 as well as H3K27Ac by Western blotting. Tri-methylation of Lysine 4 and acetylation of Lysine 27 on histone 3 are generally regarded as positive indicators of transcriptional activation, promoting gene expression (42–45). Depletion of Ambra1 or Akap8 reduced both di- and tri-methylation of H3K4 as well as acetylation of H3K27 (irrespective of whether FAK was present or not). This indicates that both Ambra1 and Akap8 can influence histone modifications, likely as a result of interacting histone modifying enzymes, which could result in chromatin remodelling and altered accessibility of transcription factors.

**Figure 6:**
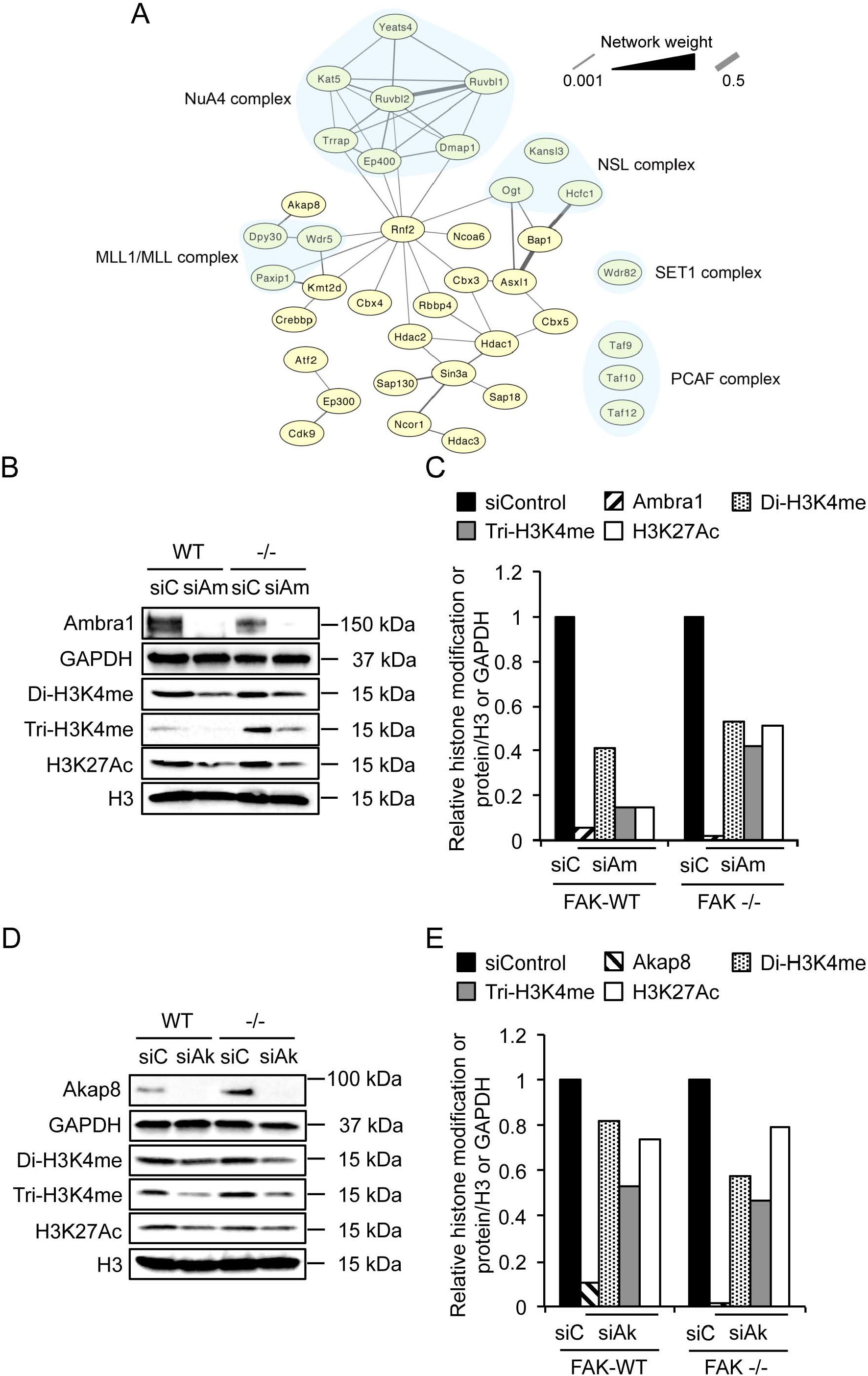
Ambra1 or Akap8 depletion decreases histone modifications. **(A)** Functional interaction network analysis of nuclear Ambra1 binding partners identified by mass spectrometry were filtered for statistical significant (p < 0.05) 2-fold enrichment over IgG control and components of histone modification complexes were subsequently used to build a protein interaction network based on direct physical interaction (grey lines). These complexes include histone acetylation complexes (NSL: Non-specific lethal); NuA4: nucleosome acetyltransferase of H4; PCAF: p300/CBP-associated factor) and histone methylation complexes (MLL1/MLL: mixed-lineage leukemia 1; SET1) (all highlighted by light blue background). **(B – E)** SCC FAK-WT and -/- cells were transfected with siControl and siAmbra1 **(B, C)** or siControl and siAkap8 respectively (siGENOME pool) (**D, E)**. After 48 h whole cell lysates were subjected to Western blot analysis using anti-Ambra1, anti-Akap8, anti-di-H3K4me, anti-tri-H3K4me and anti-H3K27Ac. Anti-GAPDH and anti-Histone H3 served as loading controls. **(C, E)** The graphs show relative histone modification levels normalised to Histone H3 or protein levels normalised to GAPDH upon Ambra1 **(C)** or Akap8 **(E)** knockdown.

## DISCUSSION

Here we describe a completely novel transcriptional signalling pathway controlled by the scaffold protein Ambra1 in the nucleus. Like Ambra1, other proteins involved in autophagy have been reported in the nucleus, e.g. LC3B binds to Lamin B1, mediating the degradation of the nuclear lamina and Beclin 1, promoting autophagy-independent DNA damage repair (46,47). No typical nuclear localisation sequence is evident for Ambra1, and hence the mechanism of nuclear translocation is unknown; however, as nuclear Ambra1 interacts with components of nuclear pore complexes and importins (Figure 1C), it is perhaps likely that nuclear import of Ambra1 occurs via binding to these in some way. Nuclear Ambra1 interacts with chromatin modifiers and transcriptional regulators in the nucleus, including those also identified as proteins that bind to FAK and IL33, e.g. SMARCC1, Ruvbl1 and Ruvbl2 (15), suggesting there may be a link between Ambra1 and FAK functions in the nucleus as well as in the cytoplasm (4). In this regard, we did find that depletion of Ambra1 leads to reduced FAK recruitment to chromatin.

The PKA-scaffold Akap8 that binds to Ambra1 in the nucleus has itself previously been linked to histone modifications and chromatin changes. Indeed, by interacting with the MLL1/MLL complex via Dpy30, Akap8 regulates histone H3K4 methyltransferase complexes and binds to the nuclear matrix, nucleoporin component Tpr as well as chromatin; in turn, this contributes to chromosome condensation and transcription, effects that are important for the mitotic checkpoint (41,48–51). Akap8 also binds the histone deacetylase HDAC3 and influences mitosis (52). Therefore, existing studies had already suggested a scaffolding function for Akap8 in assembly of chromatin modification complexes. Akap8 dissociates from chromatin and the nuclear matrix as a result of nuclear tyrosine phosphorylation, and it may have a role in regulation of chromatin structural changes (53). Finally, a recent study confirmed the scaffold function of Akap8 in organising nuclear microdomains, thereby controlling local cAMP for nuclear PKA regulation (54) – although it is not clear whether this is a chromatin-associated function of Akap8.

Nuclear envelope and nuclear pore components, like Nup153, associate with chromatin and regulate genome organisation and gene expression via nuclear pore complexes acting as scaffold platforms to allow the assembly and recruitment of transcription factors to the nuclear periphery (55,56). Therefore, Ambra1 might serve as a molecular scaffold that links chromatin to the nuclear pore complex, allowing transcription factor binding (such as Atf2) and resulting gene expression. When we looked at potential Atf2-binding sequences in the genes that were altered after depletion of Ambra1, Akap8 and Atf2, we found that (according to www.natural.salk.edu/CREB) *tgfb2* and *tgfb3* both have CRE_TATA boxes that might serve as Atf2 binding sites. Future work will examine whether Ambra1 influences the binding of p-Atf2 T71 at the promoters of these genes. Moreover, it is likely that some of the cancer-associated functions of Ambra1 in SCC cells (such as cancer cell invasion) are associated with the nuclear transcription signalling effects we report here, e.g. on TGFβ isoforms as well as its trafficking effects we reported previously (4).

Taken together, our data lead us to propose the following model (depicted in Figure 7); briefly, Ambra1 was already known to localise to autophagosomes in the cytoplasm and to focal adhesions, where it regulates the removal of ‘untethered’ tyrosine kinases via autophagy (2,4). We now show that Ambra1 also interacts with nuclear pore proteins, and locates to the nucleus where it is part of a network of inter-linked chromatin modifiers and transcriptional regulators, including a set of interacting proteins whose recruitment to chromatin is influenced by Ambra1. This includes the PKA-scaffold Akap8, the Mediator complex component Cdk9 and the transcription factor Atf2 in its active form. Moreover, Ambra1, Akap8, Cdk9 and Atf2 co-regulate the expression of a subset of genes. Both Ambra1 and Akap8 influence cellular histone modification that could contribute to their transcriptional effects. Therefore, we have uncovered a completely novel function for the autophagy protein Ambra1, which acts as a nuclear scaffold to recruit other scaffold proteins, chromatin modifiers and transcriptional regulators to elicit gene expression changes via Atf2. A similar scaffolding mechanism creating nuclear ‘transcription signalling hubs’ has been described for FAK, controlling *ccl5, IL33, tgfb2* and *igfbp3* transcription as well as for mAKAPβ, which creates nuclear ‘signalosomes’ and binds the transcriptional regulators NFAT, MEF2 and HIF1α (16,57,58).

**Figure 7:**
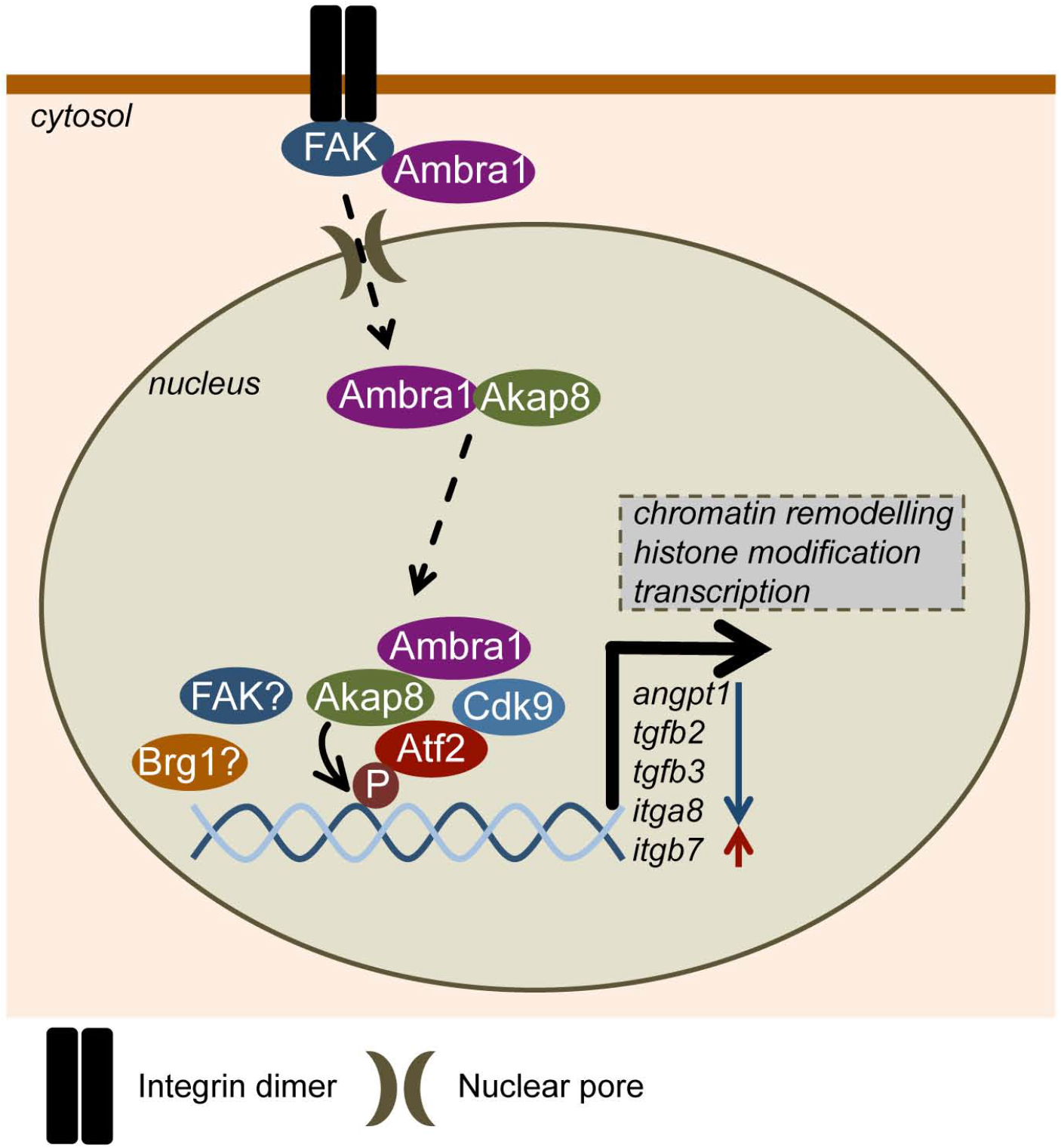
Model depicting Ambra1-dependent transcriptional regulation. In mouse SCC cells, Ambra1 is already known to localise to autophagosomes and focal adhesions, where it binds FAK and Src and regulates the removal of untethered kinases via autophagy. Ambra1 can also interact with nuclear pore components and is translocated into the nucleus most likely via nuclear pores and importins. Nuclear Ambra1 is part of a network consisting of chromatin modifiers and transcriptional regulators, some of which are recruited to chromatin in an Ambra1-mediated manner, including the PKA-scaffold Akap8, Cdk9 and active Atf2 (p-Atf2 T71). Further, Ambra1, Akap8, Cdk9 and Atf2 co-regulate the expression of a sub-set of genes like *angpt1, tgfb2, tgfb3, itga8* and *itgb7*. Both Ambra1 and Akap8 influence cellular histone modifications, which could contribute to their transcriptional effects. Overall, the autophagy protein Ambra1 also acts as a nuclear platform to recruit key scaffolds, chromatin modifiers and transcriptional regulators to elicit gene expression changes via Atf2.

## MATERIALS AND METHODS

### Antibodies

Antibodies used were as follows: anti-Paxillin and anti-GM130 antibodies (BD Transduction Laboratories, New Jersey, USA), anti-Akap8 and anti-Nup153 (Abcam, Cambridge, UK), anti-FAK, anti-PDI, anti-p-Atf2 T71, anti-Atf2, anti-Brg1, anti-Rpb1, anti-Cdk8, anti-Cdk9, anti-Histone H4, anti-H3K4me2, anti-H3K4me3, anti-H3K27Ac, anti-Histone H3, anti-Lamin A/C and anti-GAPDH (Cell Signaling Technologies, Danvers, MA, USA), as well as anti-Ambra1 antibody (Millipore, Billerica, MA, USA). Anti-rabbit or mouse peroxidase-conjugated secondary antibodies were purchased from Cell Signaling Technologies.

### Cell culture

FAK-deficient Squamous Cell Carcinoma (SCC) cell lines were generated as described previously (18). SCCs were maintained in Glasgow MEM containing 10% FCS, 2 mM L-glutamine, nonessential amino acids, sodium pyruvate and MEM vitamins at 37°C, 5% CO2. SCC FAK-WT cells were maintained in 1 mg/ml hygromycin B.

### siRNA

FAK-WT or FAK -/- SCC cells were transiently transfected using HiPerFect (Qiagen, Manchester, UK), according to manufacturer’s protocol with a final concentration of 80 or 100 nM siRNA respectively (Supplementary Material and Methods, Table 1). Cells were analyzed at 48 h post transfection.

### Whole cell lysates

Cells were washed twice in ice-cold PBS and then lysed in RIPA buffer (50 mM Tris-HCl pH 8.0, 150 mM NaCl, 1% Triton X-100, 0.1% SDS and 0.5% sodium deoxycholate) supplemented with PhosStop and Complete Ultra Protease Inhibitor tablets (Roche, Welwyn Garden City, UK) and cleared by centrifugation.

### Nuclear fractionation

Cells were washed twice in ice-cold PBS and then lysed in DET buffer (150 mM NaCl, 25 mM Hepes pH 7.5, 1 mM β-Mercaptoethanol, 0.2 mM CaCl2, 0.5 mM MgCl2, 0.5% NP-40) supplemented with PhosStop and Complete Ultra Protease Inhibitor tablets (Roche, Welwyn Garden City, UK). Lysates were incubated on ice for 10 minutes and centrifuged. The resulting pellets were washed twice in DET buffer, resuspended in RIPA buffer and cleared by centrifugation.

For mass spectrometry, nuclear lysates were prepared using Nuclei PURE prep isolation kit (Sigma, Gillingham, UK).

### Chromatin isolation

The protocol was adapted from McAndrew *et al.,* (59). All buffers were supplemented with PhosStop and Complete Ultra Protease Inhibitor tablets (Roche, Welwyn Garden City, UK). Briefly, cells were washed twice in icecold PBS, lysed in extraction buffer (10 mM Hepes pH 7.9, 10 mM KCl, 1.5 mM MgCl_2_, 0.34M sucrose, 10% glycerol, 0.2% NP-40) and centrifuged for 5 minutes at 6500 g. Nuclear pellets were washed in extraction buffer without NP-40 and centrifuged for 5 minutes at 6500 g. Pellets were resuspended in Low salt buffer (10 mM Hepes pH 7.9, 3 mM EDTA, 0.2 mM EGTA) and incubated for 30 minutes at 4 **°**C with rotation, before centrifugation for 5 minutes at 6500 g. Pellets were resuspended in High salt buffer (50 mM Tris-HCl pH 8.0, 2.5 M NaCl, 0.05% NP-40) and incubated for 30 minutes at 4 **°**C with rotation. Supernatants containing chromatin fractions were cleared by centrifugation. Proteins were precipitated by adding TCA to a final volume of 10% and incubating on ice for 15 minutes. Precipitated proteins were pelleted by centrifugation, washed twice with ice-cold acetone and then resuspended in 2x sample buffer prior to analysis by Western blotting.

### Immunoblotting, immunoprecipitation and mass spectrometry

Protein concentration was calculated using a BCA protein assay kit (Thermo Scientific, Loughborough, UK). For immunoprecipitation, 1 mg lysates were incubated with 2 μg of unconjugated antibodies at 4°C overnight with agitation. Unconjugated antibody samples were incubated with 10 μl of Protein A agarose for 1 h at 4°C. Beads were washed three times in lysis buffer and once in 0.6 M LiCl, resuspended in 20 μl 2x sample buffer (130 mM Tris pH 6.8, 20% glycerol, 5% SDS, 8% β-mercaptoethanol, bromphenol blue) and heated for 5 minutes at 95°C. Samples were then subjected to SDS-PAGE analysis as described elsewhere (4).

For mass spectrometry, Ambra1 was immunoprecipitated from 2 mg nuclear lysates of SCC FAK-WT and -/- cells (samples in triplicates), using 2 μg of unconjugated antibodies (anti-Ambra1 and rabbit-anti-IgG) at 4°C overnight with agitation. Unconjugated antibody samples were incubated with 20 μl of Protein A agarose for 1 h at 4°C. Beads were washed twice in lysis buffer and twice in PBS. Protein complexes were subjected to on-bead proteolytic digestion, followed by desalting and liquid chromatography-tandem mass spectrometry as reported previously (60). The interaction network analysis was performed using Cytoscape.

### Immunofluorescence microscopy and image analysis

Cells were fixed, stained and imaged as described in Schoenherr *et al.,* (4).

### qRT-PCR

RNA from cells was isolated using the RNeasy Mini Kit (Qiagen, Manchester, UK). 500 ng of total RNA was reverse-transcribed using the SuperScript First-Strand cDNA synthesis kit (Life Technology, Paisley, UK). For the PCR amplification in a Step One Plus real-time PCR system (Life Technology, Paisley, UK) 25 ng cDNA were used in a total reaction mix of 20 μl containing 10 ul Sensi Fast SYBR Green Hi-Rox (Bioline, London, UK) as well as 400 nM forward and reverse primer (Supplementary Material and Methods, Table 2). GAPDH was used to control for differences in cDNA input. Relative expression was calculated according to the ΔΔCt quantification method. Each sample within an experiment was performed in triplicate and the experiment was carried out three times.

### Nanostring gene expression analysis

SCC FAK-WT cells were transfected with siControl, siAmbra1, siAkap8 and siAtf2. RNA from cells was isolated 48 h post transfection using the RNease Mini Kit (Qiagen, Manchester, UK) and diluted to 20 ng/μl. Samples (in triplicates) were subjected to gene expression analysis using the PanCancer Mouse Pathways Panel (Nanostring, Amersham, UK). Analysis was performed using nSolver analysis software (Nanostring, Amersham, UK). The cut-off point of statistically significant relative changes (siRNA/siControl p < 0.05) was set to 2-fold.

### Statistical tests

For all experiments shown, *n* = 3 – 5. Error bars for the graphs show s.d. Student’s *t*-test was carried out to calculate the statistical significance.

## ACKNOWLEDGEMENTS

This work was funded by CR-UK Programme Grant (C157/A24837) to M Frame and V Brunton. We would like to thank Jimi Wills and Alexander von Kriegsheim at the IGMM Mass Spectrometry Facility for the mass spectrometry analysis and Alison Munro at the IGMM HTPU array facility for the Nanostring analysis.

## AUTHOR CONTRIBUTIONS

C.S. designed and performed the cell biology and biochemical experiments and co-wrote the paper. M.C.F. conceived proteomics, provided funding through competitively awarded grants and cowrote the paper.

